# L-type calcium channels link oxidative stress to calcium signaling pathway and membrane excitability: Insights from computational modeling of dopaminergic neurons

**DOI:** 10.1101/2022.08.16.504074

**Authors:** M.A. Andres, S. Karratti-Abordo, C. Bryan, A. Shoji, M. Zaporteza, A.M. Castelfranco

## Abstract

Dopamine neurons, which are critical in movement, cognition, and reward learning, are vulnerable to oxidative stress during aging, drug addiction and viral infection, and can lead to neurodegeneration. Previous work used computational modeling to study dopamine neuron function based on experimental findings from rodent brain slices containing dopamine neurons. Here we show the feasibility and utility of applying such computational models of DA neurons to the analysis of experimental findings from in-vitro cultured cells. DCFH-DA (and DHE) and time-lapse, Fura-2 ratiometric imaging was used to measure changes in ROS levels and changes in intracellular calcium (Ca^2+^) levels, respectively, in two dopaminergic cell models: differentiated SH-SY5Y and differentiated human neural progenitor cells. We investigated how peroxide-dependent changes in the behavior of the L-type channel might alter the excitability of the dopaminergic cell. We found that L-type channels mediated clusters of calcium spikes (or oscillations) and that our model suggested that such increased excitability could be explained by changes in the voltage-dependence of activation of the L-type channels in response to exogenous peroxide. Our findings suggested that L-type channels link oxidative stress responses to modulation of excitability, indicating that the Ca^2+^ channel blocker nicardipine may help disrupt this link by reducing oxidative stress and preventing channel activation at more hyperpolarized potentials, thus reducing plasma membrane excitability.

## INTRODUCTION

Previous studies demonstrated that oxidative stress plays a significant role in the pathology of neurodegeneration linked to Parkinson Disease (PD) [1], Alzheimer Disease [2], HIV-infection [3], and drug addiction [4]. Oxidative stress involves the generation of reactive oxygen species (ROS) like hydroxyl radicals, hypochlorite and peroxynitrites, including hydrogen peroxide (H_2_O_2_), a small molecule that can easily cross the plasma membrane and can, therefore, serve as a signaling molecule [5, 6].

In the cardiovascular system, H_2_O_2_ can modify ion channel function by increasing L-type calcium channel activity [7–9]. Hence, during oxidative stress, increases in H_2_O_2_ exposure may lead to calcium homeostasis dysfunction. It is thought that the modulation of ion transport by peroxide in signaling pathways may underlie the pathology of hypertension, cardiomyopathies, cardiac hypertrophy, congestive heart failure, and ischaemic heart disease [10]. Similarly, in the central nervous system, generation of peroxide may have deleterious effects as enhanced oxidative stress may also impair the normal functioning of dopaminergic (DA) neurons in the brain. Because ion channels are crucial in maintaining the physiology of DA neurons, any modifications to their functions due to the presence of increased ROS production may significantly affect neurons’ ability to fire action potentials, hence, modifying their excitability, impeding their communication both extracellularly and intracellularly, and disrupting the activity-dependent release of dopamine necessary for the proper functioning of downstream structures, such as the striatum [11].

Midbrain DA neurons in the substantia nigra pars compacta (SNc) are particularly vulnerable to degeneration in PD. These neurons are spontaneously active; firing action potentials in the absence of synaptic inputs [12–14]. L-type calcium channels in SNc DA neurons are active during autonomous firing, including the subthreshold interspike interval between action potentials, but they are not essential to the generation of pacemaking activity [15]. Calcium influx through these channels underlies cytosolic calcium oscillations in both the soma and dendrites [15–18]. Two-photon dendritic calcium imaging has confirmed that SNc dopaminergic neurons rely heavily on low-threshold L-type calcium channels for both tonic pacemaking and phasic calcium influx, directly linking this chronic low-threshold calcium load to accelerated metabolic stress [19]. This activity-dependent increase in intracellular calcium levels raises the metabolic costs associated with calcium homeostasis since calcium must be extruded by ATP-dependent processes resulting in increased oxidative stress in the mitochondria and endoplasmic reticulum (ER), and production of superoxide and other reactive oxygen species [20–22]. The prevailing model is that L-type channels mediate the Ca^2+^ influx that drives the sequestration of Ca^2+^ ions into the ER and subsequently into the mitochondria, where sustained Ca^2+^ levels lead to the stimulation of mitochondrial nitric oxide synthase, which enhances the production of ROS, hence oxidative stress [23]. Recent research suggests that perturbations in calcium microdomains serve as critical upstream triggers that shift calcium from an active signaling mechanism to a major driver of neurodegenerative pathways, including cross-talk with iron homeostasis and cell death pathways involving ferroptosis [24]. Over time, aging and genetic impairment of mitochondrial function can lead to degeneration of DA neurons. In addition to this autonomous mitrochondrial oxidative stress, there is evidence that external sources of oxidative stress from inflammation and production of ROS and reactive nitrogen species could accelerate the loss of SNc DA neurons and the progression of PD [25]. In this study, we focused on the effects of one such external source of oxidative stress, exogenous H_2_O_2._ We explored the notion that there could be positive feedback in which H_2_O_2_ might serve as a mediator molecule because of its ability to directly regulate L-type channel activity, which can,in turn, influence pacemaking.

Human DA neurons are difficult to obtain and maintain in primary culture, so numerious studies have used the SH-SY5Y cell line to investigate the pathogenesis of PD (Xicoy et al., 2017). The neuroblastoma dervived SH-SY5Y cells exhibit catecholaminergic neuronal properties but are not purely dopaminergic, although they express tyrosine hydroxylase - a marker used to identify dopaminergic phenotype [26–31]. Their oncogenic properties raise questions about their suitability as PD models. Another approach is to use differentiated human neural progenitor cells (hNPCs) as an *in vitro* human cell culture system to study neurodegenerative diseases.

Differentiated hNPCs are capable of developing a functional neuronal phenotype including generating action potentials (Young et al., 2011). They express a range of voltage-dependent ion channels including a variety of sodium, calcium and potassium channel subunits (Young et al., 2011). hNPCS express tyrosine hydroxylase following differentiation indicating that they are dopaminergic [32, 33].

This study investigated changes that exogenous H2O2 may exert on calcium signaling and excitability in two DA cell models: differentiated SH-SY5Y and differentiated hNPCs, and explored the mechanisms of action of H2O2 using a computational DA neuron model.

Biophysical cellular models are increasingly used as a basis for multiscale computational modeling in Parkinson’s disease, linking mathematical channel kinetics to mitochondrial energetic needs, dopamine synthesis, and systemic cellular death [34]. We hypothesized that during enhanced oxidative stress, peroxide might trigger and reshape action potentials and overall membrane excitability; it might do so via a direct effect on L-type calcium channel behavior and function.

## EXPERIMENTAL PROCEDURES

### Culturing SH-SY5Y cells

Adherent and undifferentiated SH-SY5Y cells were grown under 37°C and 5% CO_2_ conditions in a medium consisting of DMEM (Mediatech Inc., VA), 10% fetal bovine serum (HyClone Laboratories Inc., UT), 2mM L-alanyl-L-glutamine (HyClone Laboratories Inc., UT), and 1x Penicillin-Streptomycin (Thermo Fisher Scientific, MA). Cells were plated on poly-L-ornithine (0.01% solution; Sigma-Aldrich Co., MO) or poly-L-lysine (0.01% solution; Sigma-Aldrich Co., MO)-coated Petri-dishes. Then, the cells were differentiated using 10µM all-trans retinoic acid (ATRA) in supplemented DMEM medium. Fifty percent of the cell medium was removed and replaced with fresh ATRA supplemented medium. Differentiating medium was then replaced every 3-4 days.

### Culturing hNPCs

Human neural progenitor cells (hNPCs) (Neuromics, MN) were plated on Matrigel (Corning, NY)-coated glass-bottom Petri-dishes. hNPCs were maintained in growth medium, consisting of AB2 Basal Neural Media (ArunA Biomedical, GA), ANS Neural Supplement (ArunA Biomedical, GA), 1x Penicillin-Streptomycin (Gibco, MD), 2mM L-alanyl-L-glutamine (HyClone Laboratories Inc., UT), 1X B-27 supplement (Thermo Fisher Scientific, MA), 10 ng/mL recombinant human leukemia inhibitory factor (LIF; PeproTech US, NJ), and 20ng/mL recombinant human fibroblast growth factor-basic (FGF-b; PeproTech US, NJ). hNPCs were maintained under 37°C and 5% CO_2_ conditions. Media were replaced every 3-4 days, and cells were grown to ∼100% confluence before passaging. hNPCs were plated on poly-L-ornithine-and laminin-coated plates and were differentiated with growth media (without FGF-b) supplemented with 25ng/mL recombinant human glial-derived neurotrophic factor (GDNF; PeproTech US, NJ). Fifty percent of cell media were removed every three to four days and replaced with fresh GDNF supplemented media. Cells were differentiated for either 4-7 days before conducting time-lapse Fura-2AM ratiometric calcium imaging experiments, or 7 or more days before performing ROS assays.

### ROS assay

Differentiated SH-SY5Y and hNPCs cells were washed with unsupplemented (plain) neurobasal media (Thermo Fisher Scientific, MA) and treated with H_2_O_2_ (SH-SY5Y cells: 1mMH_2_O_2_; hNPCs: 250 µM H_2_O_2_) (and other agents as indicated in the text) for 30 minutes. Cells were then washed three times with unsupplemented neurobasal media and loaded with 200µM 2′,7′-dichlorofluorescein diacetate (DCFH-DA) fluorescent dye (Thermo Fisher Scientific, MA) for 30 minutes at 37°C. Loaded cells were washed three times again with plain neurobasal media to remove any residual DCFH-DA dye. Then, treated cells were lysed with 111µl of 90% DMSO/10%PBS for 10 minutes at room temperature in the dark, shaking on a rocker to facilitate detachment from the plate. One hundred microliters of lysed cells in the DMSO solution were transferred to wells in black 96-well plates. The plate was read within 1 hour after cell lysis using a SpectraMax plate reader (Molecular Devices, CA). Samples were excited at 480nm, and fluorescent intensity was read at 530nm emission.

Intracellular ROS levels were also detected with a fluorescent microscope using the fluorescent dye, dihydroethidium (DHE), otherwise known as hydroethidine. Cultured, differentiated SH-SY5Y cells, plated on glass-bottom petri dishes, were loaded with 35 µM DHE for an average of 30 minutes at 37°C. Then, time-lapse images of fluorescent intensities were acquired as cells were perfused first under control (see below) and then under different drug conditions or high potassium Tyrode solution consisting of (in mM): 5 NaCl, 129 KCl, 2 CaCl_2_, 1 MgCl_2_, 30 Glucose, 25 HEPES. Loaded cells were excited at 480nm and fluorescent images were captured at 590nm emission. For the analysis, after background subtraction, ROS levels (as indicated by the ratio F/F_0_) were calculated by normalizing the fluorescence intensities (F) under drug conditions to the average fluorescence intensities under control saline conditions (the baseline F_0_).

### Fura-2 ratiometric Ca^2+^ measurements

Briefly, differentiated hNPCs and SH-SY5Y, plated on coated-glass bottom dishes, were loaded with 1 µM Fura-2AM fluorescent dye (Thermo Fisher Scientific, MA) for 15 minutes in the dark at room temperature. Cells were then placed on the microscope stage and perfused with the control Tyrode solution consisting of (in mM): 129 NaCl, 5 KCl, 2 CaCl_2_, 1 MgCl_2_, 30 glucose, and 25 HEPES, then followed by perfusion of Tyrode solution containing H_2_O_2_ or the drugs (as indicated in the text). A complete experiment consisted of 3-4 minutes in normal saline (Tyrode solution) followed by 20 minutes perfusion of peroxide alone or with nicardipine in Tyrode solution. Cells were then excited at 340 and 380 nm wavelengths at 10-second intervals. The light emitted above 490 nm was captured using an intensified CCD camera interfaced with the inverted microscope. After background subtraction, Ca^2+^ levels were expressed as the ratio of fluorescence intensity excited at 340/380 nm. Time-lapse measurements of fluorescent intensities from cell images were acquired and analyzed using Winfluor Software (University of Strathclyde, Glasgow, Scotland). Averaged 340/380 ratios were calculated from imaged cells/ regions of interest (ROIs).

### Immunocytochemistry

Cells were first fixed with 2% paraformaldehyde for 10 minutes at room temperature then washed three times with 1x PBS. Then, we permeabilized the cells with 1x PBST for 10 minutes at room temperature and blocked with 1% normal goat serum in PBST for 10 minutes at room temperature. Next, we washed the cells with PBST three more times. Cells were then incubated with a 1:100 dilution of rabbit polyclonal antibody against tyrosine hydroxylase (TH (H-196), Santa Cruz Biotechnology Inc, TX) and a 1:200 dilution of antibody against dopamine transporter (DAT, Millipore Sigma, MA) for 1 hour at room temperature and washed three times before incubating with 1:1,000 dilution of goat anti-rabbit secondary antibody against Alexa Fluor 488 (Thermo Fisher Scientific, MA) for 40 minutes at room temperature. Cells were washed three times with 1x PBST and mounted with DAPI Fluoromount-G^®^ (SouthernBiotech, AL).

### Chemical Compounds

All the chemicals used for this study are at least 95% pure as determined by methods specified at the manufacturer’s website. Chemicals listed for Tyrode solutions were purchased from Sigma.

### Statistical analysis

Using ANOVA, we compared the ROS levels of the different treatments to the levels under control saline solution/media. Post-hoc comparisons were conducted using Tukey’s HSD test. ROS levels were presented as mean values ± SEM. The p-value *p*<0.05 was considered significant.

### Computational modeling

Model DA neurons were simulated using the NEURON simulation environment (version 7.1) [35, 36]. The DA neuron model was modified from that of Chan, Guzman [16] available from the modelDB database (https://modeldb.science/97860), with the modeled ion channel densities adjusted to the levels given in Guzman, Sanchez-Padilla [15]. To accurately replicate the amplitude and duration of action potentials of DA neurons, the kinetic description of the fast sodium current was based on physiological measurements of Seutin and Engel [37]. This sodium current was described using the Hodgkin-Huxley formalism [38]:

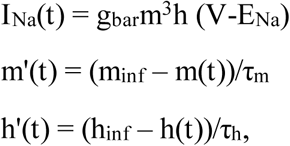

with voltage-dependent steady-state activation and inactivation functions given by:

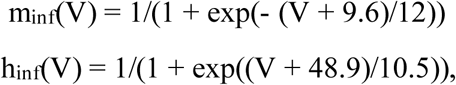

and the time constants by:

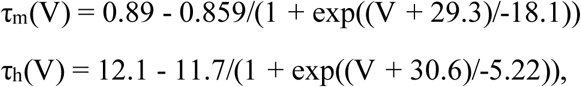

where I_Na_(t) is the sodium current at time t, m(t) and h(t) are activation and inactivation, respectively, and V is the membrane potential. The maximum conductance of I_Na_ was adjusted to match recorded spike amplitudes giving g_bar_ = 0.05 S/cm^2^ and the sodium reversal potential was set to E_Na_ = 50 mV [15]. The resulting model contained the following ion channels given with their channel densities (S/cm^2^): a transient sodium channel (0.05); 4 voltage-dependent potassium channels: K_v_1 (3×10^-5^), K_v_2 (2.5×10^-3^), K_v_4 (8.5×10^-5^), K_v_7 (7×10^-7^)); 2 calcium-activated potassium channels: SK (1×10^-5^) and BK (1×10^-3^); a hyperpolarization-activated cation channel: HCN (1.25×10^-4^); 2 calcium channels: L-type (5×10^-5^) and P/Q-type (1×10^-5^), and leak (3.5×10^-6^). The model included mechanisms for intracellular Ca^2+^ diffusion, buffering and extrusion taken from the model of Chan et al. (2007), which were based on mechanisms that are bundled with NEURON (Carnevale and Hines, 2006). However, the Ca^2+^ pump current was not included in the model. The SK channel in the model is activated by calcium entry via the L–type channel during spontaneous firing, and blocking the SK channel results in increased firing frequency. The morphology of the model neuron was based on measurements of differentiated hNPCs and consisted of a cylindrical soma, 4.1 µm in diameter and 6.3 µm in length, with three thin cylindrical processes, each having a 0.72 µm diameter and 80 µm length. The cytoplasmic resistivity was set to 70 Ωcm; the specific membrane capacitance to 1µF/cm^2^, and the reversal potential for the leak current was-20 mV. Ion channel densities in both the soma and the processes were set to the same values. To model the effects of H_2_O_2_ on the L-type calcium channel the voltage-dependence of the steady-state activation function,

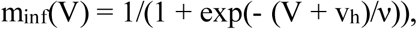

was shifted by changing the half-activation (v_h_) and/or the slope (ν) parameter. The kinetic description of L-type calcium channel includes calcium dependent-inactivation but no voltage-dependent inactivation mechanism [16].

The simulations used NEURON’s default backward Euler integration method with a time step-size of 2.5 µs. The soma and processes were subdivided into 2 µm and 27 µm compartments, respectively. A 3-fold increase in the fineness of the spatial grid resulted in less than a 0.1% change in the spontaneous firing frequency.

## RESULTS

In this study, we sought to investigate whether a relationship between oxidative stress and calcium signaling exists in dopaminergic cells as it does in cardiac cells. We used differentiated SH-SY5Y cells and differentiated hNPCs as models of DA neurons. Many neuroscience studies have used the SH-SY5Y cells as a cell model for DA neurons, as these cells express tyrosine hydroxylase - a marker used to identify dopaminergic phenotype [26–29, 31]. Differentiated hNPCs have also been reported to express tyrosine hydroxylase [32, 33]. Figure 1 shows that differentiated hNPCs and SH-SY5Y expressed both dopamine transporters and tyrosine hydroxylase, thus demonstrating that both have dopaminergic phenotypes. Tyrosine hydroxylase was not expressed in undifferentiated hNPCs (data not shown).

**Fig 1.**
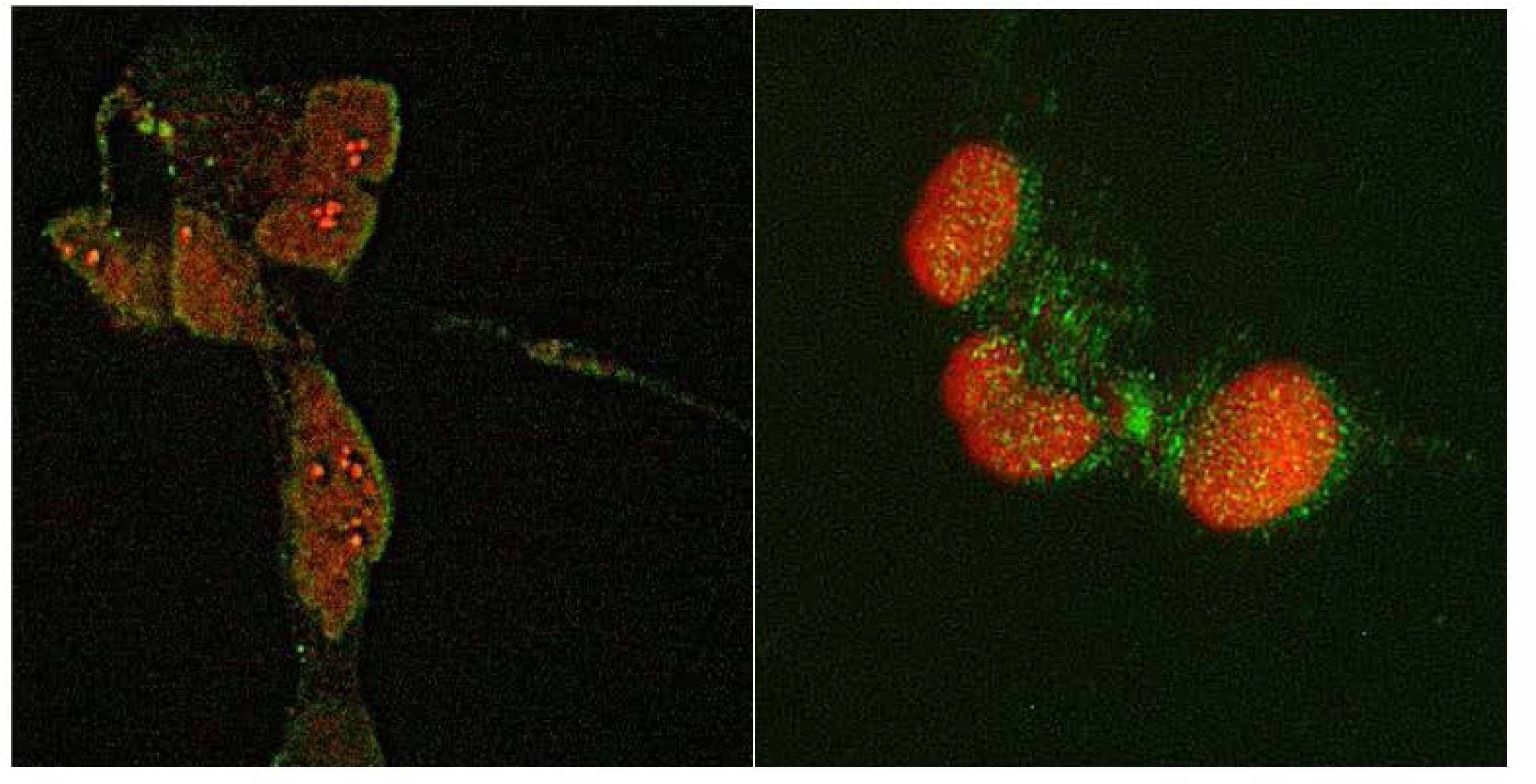
Seven-day differentiated hNPCs (left panel) and SH-SY5Y (right panel) cells were processed for immunocytochemistry. As shown, both cells are immune-positive for tyrosine hydroxylase (green), and dopamine transporter (red).

A previous patch clamp study of cardiac myocytes found that H_2_O_2_ mobilized entry of calcium ions into the cells via voltage-gated calcium channels [7], suggesting a potential role of these channels in oxidative stress. We investigated whether this role is also conserved in brain cells. We found that exogenous application of H_2_O_2_ elevated intracellular ROS levels in differentiated SH-SY5Y when compared to those cells in control media (Figure 2A). Then, we asked whether L-type Ca^2+^ channels may be involved in mediating the elevation of ROS levels. We found that L-type channel blocker, nicardipine, attenuated the ROS level increases induced by H_2_O_2_ in these dopaminergic-like cells. In particular, H_2_O_2_ in the presence of 1µM nicardipine reduced intracellular ROS to levels comparable to controls (Figure 2A). Exposure to nicardipine, with and without peroxide, did not affect ROS levels, and the cells’ responses to nicardipine are similar to those in the control media. We ascertained whether the non-cancerous, human dopaminergic neurons differentiated from hNPCs have similar oxidative stress responses to the dopaminergic model SH-SY5Y cell line derived from neuroblastoma. Figure 2 shows that dopaminergic hNPCs and SH-SY5Y cells have similar responses to H_2_O_2_ (250 µM or 1,000 µM, respectively) in that exogenous peroxide elevated ROS levels in both cells. Treatment with nicardipine (1-2 µM) with and without peroxide kept ROS at baseline levels. The antioxidant ascorbic acid (in the form of sodium ascorbate) also kept ROS at baseline levels in the presence and absence of peroxide. Together, these observations showed that not only did both the SH-SY5Y cell line and the differentiated hNPCs exhibit dopaminergic phenotype they also demonstrated similar ROS responses to exogenous application of H_2_O_2_.

**Fig 2.**
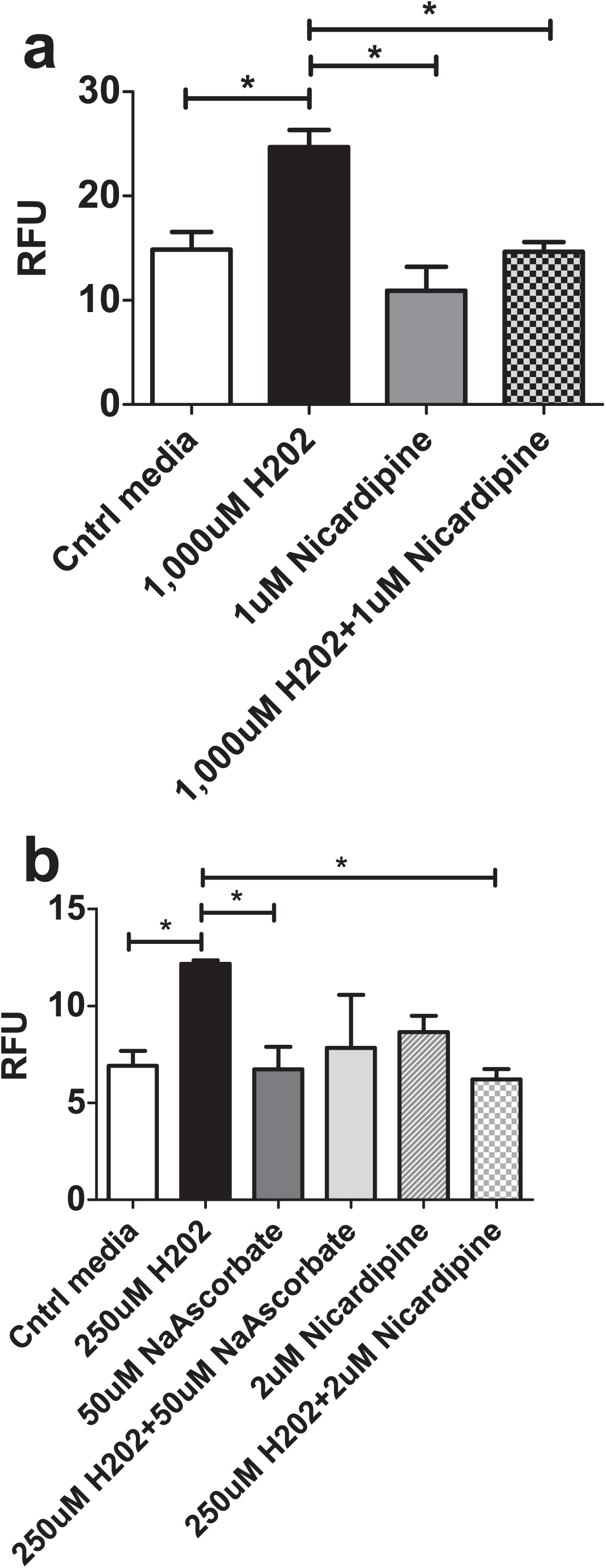
ROS levels were elevated by H_2_O_2_ and reduced to control levels by nicardipine in two models of dopaminergic neurons: (A) differentiated SH-SY5Y and (B) differentiated hNPC. (B) Sodium ascorbate, a bioavailable form of ascorbic acid, was also able to reduce the effect of peroxide to control levels in differentiated hNPC. Measurements of ROS levels were conducted using DCFH-DA fluorescent dye (each group n=3). Results are measured as relative fluorescent unit (RFU) values, which are proportional to ROS levels. Statistical significance with p<0.05 is indicated by asterisk and was determined using ANOVA with Tukey’s *post hoc* test.

Since the L-type channel blocker nicardipine was effective in reducing ROS levels, we investigated a potential mechanism for the impact of exogenous peroxide on ROS through calcium signaling using time-lapse, Fura-2 ratiometric measurements. First, we superfused dopaminergic hNPCs with control saline solution for 3-4 mins then followed with saline solution supplemented with 250µM H_2_O_2_ for 20 min. Figure 3A shows that H_2_O_2_ induced clusters of intracellular calcium increases (oscillations), increasing in amplitude with longer exposure. To determine whether these calcium increases were attributed to calcium entry via the L-type calcium channel, we exposed the dopaminergic cells with 2µM nicardipine in the presence of 250µM H_2_O_2_. We found that this treatment produced a small “bump” but prevented any large increases and oscillations in the intracellular calcium concentrations (Figures 3B and 3D).

**Fig 3.**
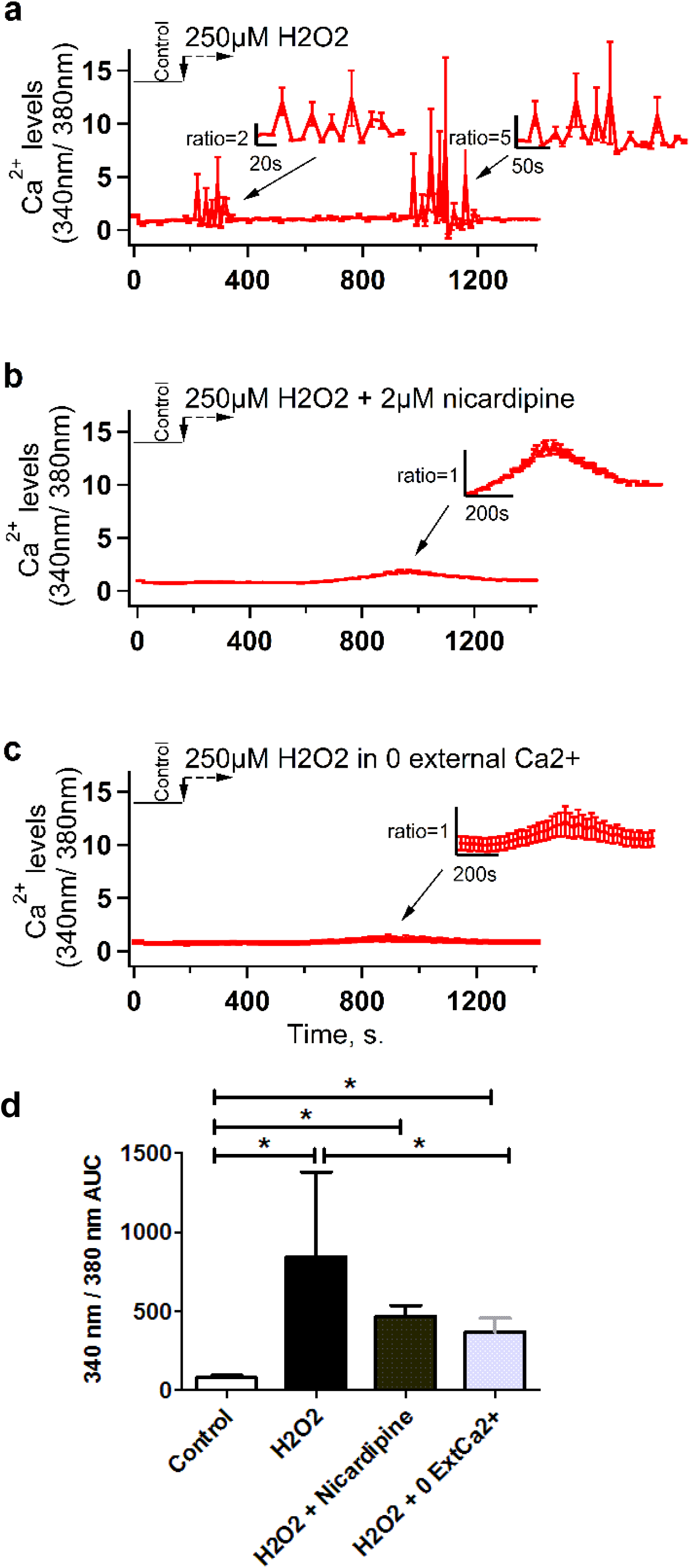
H_2_O_2_ (A) elevates intracellular Ca^2+^ levels, as expressed by the 340nm / 380nm fluorescent intensity ratio, but the presence of nicardipine (B) suppressed calcium responses (each group, n≥12 hNPCs). (C) The omission of external Ca^2+^ prevented the clusters of Ca^2+^ increases or Ca^2+^ oscillations. (D) Bar graph summarizes the area under the curve (AUC) of the cells’ responses to the various treatments in Figures 3A, 3B, and 3C. Amplified regions (designated by arrows) are shown as insets in Figures 3A-C. Asterisk indicates statistical significance with p<0.05.

Similarly, when we omitted external calcium ions in the saline solution in the presence of H_2_O_2_, there was a lack of response (oscillation) that was similar to those cells treated with nicardipine (Figure 3C and 3D). The slight bump in intracellular calcium concentration in Figures 3B and 3C suggest that intracellular stores might compensate when L-type channels are blocked by nicardipine or when extracellular calcium is omitted, suggesting that release from stores may play a more significant role when calcium entry through the plasma membrane is suppressed.

Does the increase in calcium concentration inside the cell via the L-type calcium channels lead to enhancement of ROS levels? Figure 4A demonstrates that activating L-type channels with 2µM mefenamic acid, an L-type channel agonist [39], in the absence of exogenous peroxide increased ROS levels, which can be reversed by nicardipine. In single-cell assays, using Fura-2 calcium measurements with microscopy, we found that mefenamic acid, without an oxidant, induced intracellular Ca^2+^ increases (Figure 4B, *left*). As a positive control, high potassium was used to stimulate the cells to induce Ca^2+^ responses. The presence of nicardipine (2µM) and mefenamic acid (2µM) suppressed these Ca^2+^ effects (Figure 4B, *right*). Similarly, we also observed increases in ROS levels in response to mefenamic acid (Figure 4C, left), which were also suppressed by nicardipine (Figure 4C, *right*). Together, these data suggest that the L-type calcium channels mediate H_2_O_2_-associated increases in cytosolic calcium levels and that these channels might potentially explain the H_2_O_2_-induced calcium oscillation responses.

**Fig 4.**
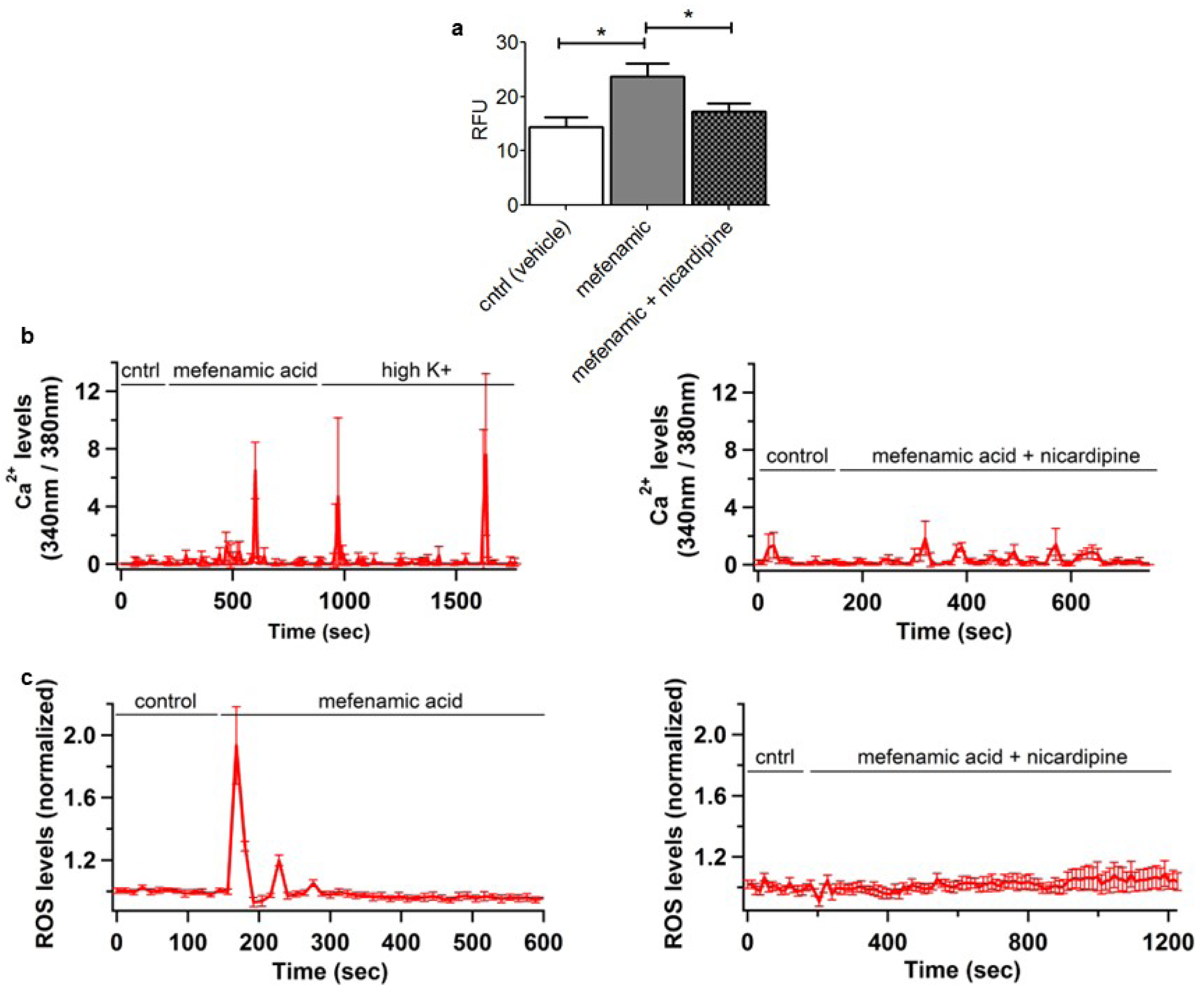
L-type channel agonist mefenamic acid enhances both ROS and Ca^2+^ responses. (A) ROS levels in DCFH-DA-loaded hNPC neurons were elevated by 2µM mefenamic and reduced to control levels by 2µM nicardipine (each group n≥4), as measured from a population of cells using a fluorometric microplate reader. Asterisk indicates statistical significance with p<0.05. (B) Fura-2 ratiometric imaging measurements on differentiated SH-SY5Y cells show increases in Ca^2+^ levels in response to mefenamic acid (2µM) and positive control high potassium Tyrode solution (left panel; n=13 ROIs). Nicardipine (2µM) in the presence of mefenamic prevented rises in intracellular Ca^2+^ (right panel; n=27). (C) Mefenamic acid (2µM) also induced ROS responses (measured as F/F_0_) in differentiated SH-SY5Y cells (left panel; n=31 ROIs). Nicardipine (2µM), together with mefenamic acid, suppressed such responses (right panel; n=15 ROIs). The fluorogenic probe DHE was used in Figure C above to detect ROS products.

We used computational modeling approaches based on models of midbrain DA neurons to investigate how hypothesized changes in L-type Ca^2+^ channel activity induced by high peroxide might lead to changes in calcium levels, as observed in the first cluster of Ca^2+^ oscillations (Figure 3A), and affect the excitability of these DA neuron-like cells. As shown in Figure 5A, the model DA neuron fired spontaneously at the rate of 3.4 Hz, which is in the physiological range of 1-8 Hz [12, 13]. This spontaneous activity was dependent on the fast, transient sodium current. Blocking the sodium channels in the model resulted in the cession of spontaneous firing, in contrast, when the L-type Ca^2+^ channels were blocked the model neuron continued to fire, although at a lower frequency (see Figure 6A). The regular firing pattern is consistent with the lack of synaptic input in the model [14, 18]. Facilitation of the L-type Ca^2+^channel by H_2_O_2_, reported in Yang, Xu [7] was modeled as a change in the voltage-dependence of the steady-state activation curve either as a shift to the left in the half-activation voltage (Figure 5B; dashed, red curve) or as a combination of a shift in the half-activation and a change in the slope (Figure 5B; dotted, blue curve). Both of these changes result in greater activation of the L-type Ca^2+^ current during the interspike interval, as shown by voltage-clamp Ca^2+^ currents for voltage steps to voltages in this range:-60 mV and-40 mV (Figure 5C, left and center panels), and in changes in the shape and period of the spontaneous action potentials (Figure 5D). This leads to an increase in the spontaneous firing frequency and the average internal calcium concentration in the soma of the model neuron relative to the default model, model 1 (Figure 6). If instead the L-type Ca^2+^ channels were blocked, the spontaneous firing frequency was reduced to 2.7 Hz, and the average calcium concentration decreased from 0.14 µM to.06 µM (Figure 6; model 4). We further investigated the effects of an increase in L-type Ca^2+^channel density such as might result from increased production of L-type calcium channel transcripts and channel proteins. Again both the spontaneous firing frequency and the internal Ca^2+^ concentration of the model cell were increased with relatively larger increases in the models with the facilitated L-type Ca^2+^ channels (Figure 6).

**Fig 5.**
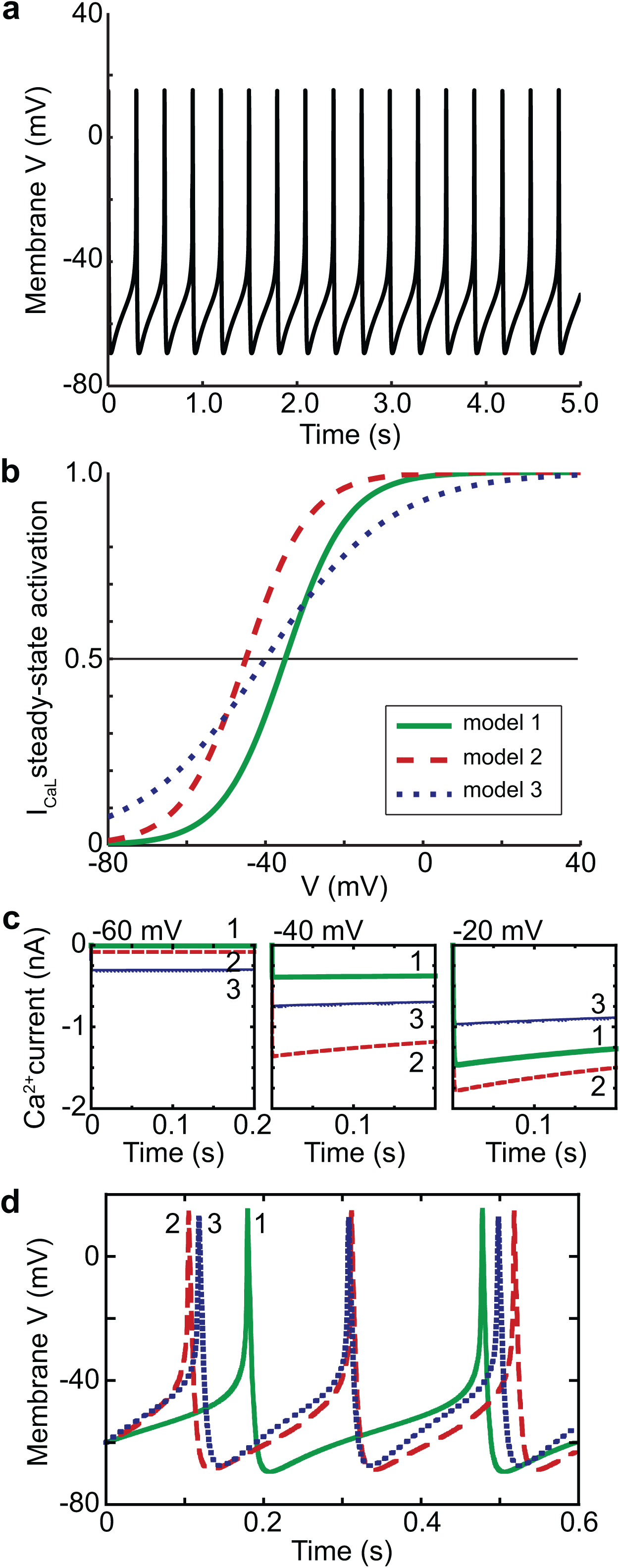
Computational modeling of DA neurons. (A) Model DA neurons fire spontaneously at 3.4 Hz. (B) Voltage-dependence of steady-state activation curve for model L-type calcium channel is shifted by H_2_O_2_: model 1 (control): v_h_ =-35 mV, ν = 8 mV(green, solid line); model 2: v_h_ =-45 mV, ν = 8 mV (red, dashed line); model 3: v_h_ =-40 mV, ν = 20 mV(blue, dotted line). (C) Simulated voltage–clamp records of L-type Ca^2+^current with control steady-state activation (model 1, green line) and activation shifted by H_2_O_2_ (model 2, red dashed line; model 3, blue line) for 3 voltage steps from-70 mV to-60 mV,-40 mV,-20 mV. Note that for each command voltage, the relative amounts of Ca^2+^ current for the models change. (D) Action potentials for the three DA neuron models shown for two cycles of model 1 (green, solid line).

**Fig 6.**
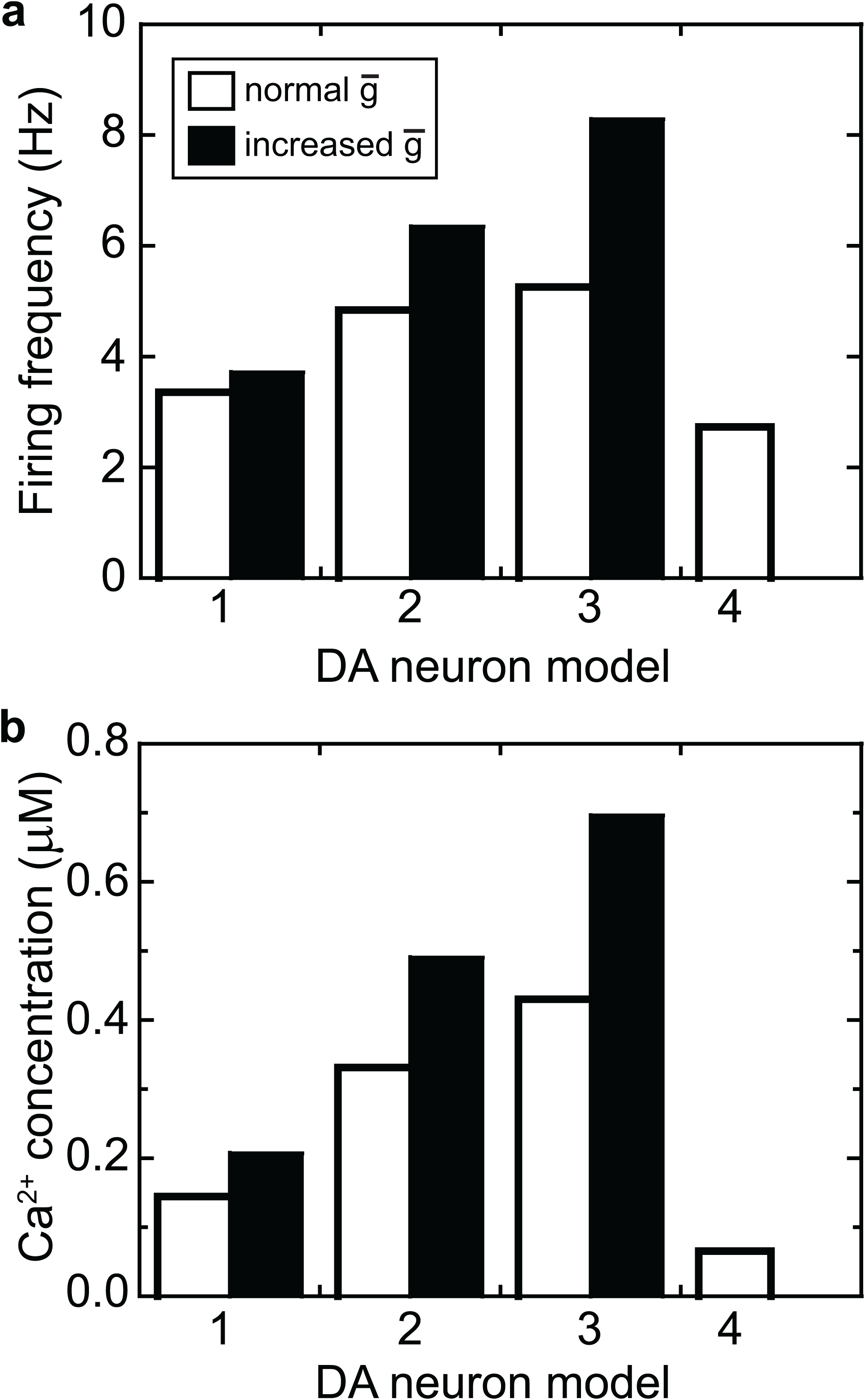
Firing frequency and intracellular Ca^2+^ concentration for DA neuron models with different L-type Ca^2+^ channels. (A) Firing frequency of DA neuron models. Models: (1) control: v_h_ =-35 mV, ν = 8 mV; (2) shifted, v_h_ =-45 mV, ν = 8 mV; (3) shifted, v_h_ =-40 mV, ν = 20 mV; (4) L-type Ca^2+^ channel blocked. Filled bars indicate models with doubled L-type Ca^2+^ channel density (g_bar_) in addition to the shift in the voltage-dependence of activation. (B) Intracellular Ca^2+^ concentration for DA neuron models. Same models as in panel a; filled bars indicate models with doubled L-type Ca^2+^channel density.

## DISCUSSION

An established mechanism is that the L-type Ca^2+^ channel in DA neurons mediates a unidirectional process linking calcium signals to mitochondrial oxidative stress [20, 23]. Questions remain as to whether L-type Ca^2+^ channel activity and behavior may provide a link in the reverse direction from oxidative stress to calcium signaling, and subsequently, mediate changes in membrane excitability. In the current study, through a combination of fluorometric microplate and microscopy assays we measured intracellular ROS (hydroxyl radicals [40] and superoxide [41]) as estimated from the fluorescent signal of DCFH-DA and DHE, respective, and used Fura-2AM ratiometric Ca^2+^ imaging techniques to measure Ca^2+^ levels, in two human dopaminergic cell models: SH-SY5Y and neurons differentiated from hNPCs. We found that exogenously applied peroxide increased intracellular ROS and Ca^2+^ levels. Time-lapse measurements showed oscillations in internal calcium levels with exposure to peroxide that may contribute to increased excitability of the differentiated hNPCs. We then used a well-established model of a DA neuron to determine whether changes in the L-type Ca^2+^ channel such as those reported for cardiac cells could explain the observed changes in the cultured DA cells [42, 43].

For the current study, the model was modified to match the morphology of the differentiated hNPCs and contained a mechanism for the direct facilitation of L-type Ca^2+^ channels on exposure to H_2_O_2_ [7]. These changes in the steady-state activation of the L-type Ca^2+^ channel resulted in increases in internal Ca^2+^ concentration. In particular, the average Ca^2+^ concentration in the facilitated models was 2-3 times that of the control model (Figure 6B). This is comparable to the increase in mean intracellular Ca^2+^ concentration, as determined by the area under the fluorescence curve, that we observed in the differentiated hNPCs exposed to H_2_O_2_ relative to cells exposed to H_2_O_2_ and nicardipine combined (see Figure 3D). Hence, even these relatively small shifts in the activation characteristics of the L-type Ca^2+^ channel could explain the increase in Ca^2+^ concentration seen in the differentiated hNPCs exposed to H_2_O_2_.

Our computational model predicted that increases in intracellular Ca^2+^ concentration mediated by enhanced L-type Ca^2+^ channel activity would increase the firing frequency of dopaminergic neurons. This occurred even though the model contained several calcium-dependent mechanisms that act to reduce excitability in response to elevated intracellular calcium: two types of calcium-dependent potassium channels (SK and BK) and calcium-dependent inactivation of the L-type Ca^2+^ channel. It is consistent with a model tuned to capture the participation of calcium channels in the pacemaking mechanism of SNc dopamine neurons that the negative feedback of SK and BK channels does not inhibit the spontaneous activity [16–18]. Calcium entry through L-type Ca^2+^ channels during pacemaker activity has been shown to lead to mitochondrial oxidative stress in SNc DA neurons [20, 21, 23]. Our study showed that exogenous H_2_O_2_ increased calcium influx through L-type Ca^2+^ channels and increased the production of ROS. Furthermore, activation of these L-type channels by the agonist, mefenamic acid, also led to a rise in ROS. This suggests positive feedback exists between exogenous H_2_O_2_, increased calcium influx through L-type Ca^2+^ channels, elevated intracellular calcium and increased mitochondrial production of ROS, including H_2_O_2_ (Figure 7). We hypothesize that this positive feedback mechanism (Figure 7) could be driven by increased calcium influx through channels potentiated by exogenous peroxide, as suggested by our calcium imaging data (Figures 3 and 4), as well as by intracellular peroxide generated by mitochondria in response to the increased calcium [7, 10, 20, 23]. Hydrogen peroxide is a small molecule that readily passes through the plasma membrane and can act through direct oxidation of the L-type Ca^2+^ channel protein, possibly at the cysteine residues [7, 10]. This also raises the question as to whether L-type channel blockers may be an appropriate therapeutic option for minimizing or protecting against further damages due to oxidative stress in dopaminergic neurons and for restoring the excitability and calcium levels of these neurons close to their normal, unperturbed state. This is significant as there are several FDA approved L-type channel blockers with known safety records that can be repurposed for protecting and preserving dopaminergic neurons from ROS-induced degeneration. One such channel blocker, isradipine, was evaluated for its efficacy in a randomized clinical trial, but was not found to slow the progression of early-stage PD relative to the placebo [44–46]. A possible limitation of the clinical trial was that the isradipine dose may have been too low to provide the necessary inhibition of the L-type Ca^2+^ channels.

**Fig 7.**
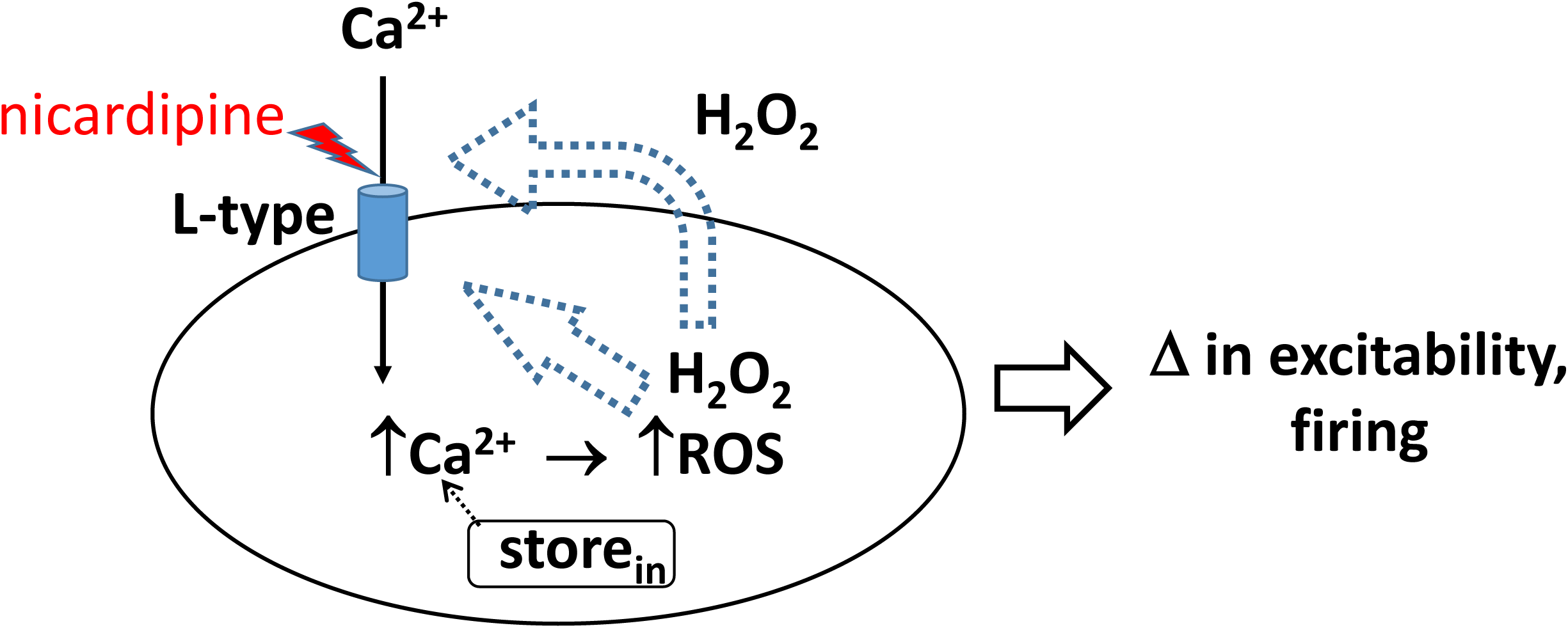
Diagram of the proposed positive feedback loop. Positive feedback occurs when peroxide, generated internally during oxidative stress, acts directly on the L-type channels. Because it can easily cross the plasma membrane, peroxide can also diffuse out of neurons and glial cells, during neuro-inflammatory responses, and act on the L-type channels extracellularly. Both exogenous and endogenous peroxide facilitates calcium influx via L-type calcium channels, which then leads to further increases in the production of reactive oxygen species, including peroxide. The application of L-type channel blockers such as nicardipine could disrupt this feedback.

Our time-lapse measurements found two events of increased intracellular calcium oscillations with exposure to H_2_O_2_ separated by a relatively quiescent period of about 10 minutes, with the second event showing increased peak calcium levels. It is possible that these three periods (i.e., oscillation, quiescent, oscillation) indicated a shift in underlying, secondary processes in response to H_2_O_2_. The intracellular calcium measurements reflected changes in fluorescence of a small group of hNPCs sampled at 10-second intervals, so the observed frequency in the oscillations was dependent on the sampling rate. Increased calcium levels could result in part from increased overlap in the peak fluorescence of the individual cells. This could be due to increased firing frequency and possibly synchrony in the firing of the cells in the dish due to synaptic connections or a shift to firing prolonged bursts of action potentials. Our model predicted that shifts in the activation properties of the L-type Ca^2+^ channel could result in increased firing frequencies and calcium influx and that this would be further enhanced by a mechanism that leads to increased expression of L-type channels. However, the changes in firing frequency predicted by the model were on a much faster time scale, and hence, are difficult to translate to the time scale of the observed oscillations without incorporating additional mechanisms. Other redox-sensitive channels such as the ATP-sensitive potassium (K_ATP_) channel or the A-type Kv4.3 potassium channel could be involved in the response to exogenous H_2_O_2_ possibly underlying the quiescent phase. Changes in the activation of these channels might produce greater changes in firing properties of the hNPCs including a shift to bursting, which might lead to the second cluster of Ca^2+^ oscillations [47–50]. Additionally, ryanodine receptors (RyR), known to be localized in the ER, play an essential role in the intracellular Ca^2+^ signaling, as their blockage leads to the reduction of cytosolic Ca^2+^ during pacemaking [23]. We speculate that the second cluster of oscillations may induce the activation of RYRs in the ER to amplify the Ca^2+^ signal of the cells’ response to H_2_O_2_. It is worth noting that the RYR location in the ER also creates a mechanism by which Ca^2+^ can easily be moved from the ER to the mitochondria, where sustained levels of Ca^2+^ can drive the production of ROS via stimulation of nitric oxide synthase [23].

In addition to exhibiting a dopaminergic phenotype, both the SH-SY5Y cells and the differentiated hNPCs demonstrate similar ROS responses to exogenous H_2_O_2_. They also share similarities (as well as differences) in the expression of calcium channels: while SH-SY5Y cells express both the L-type Cav1.2 (encoded by *CACNA1C* gene) and Cav1.3 (*CACNA1D* gene) channel subtypes [51], differentiated hNPCs express the *CACNA1C* gene that encodes the Cav1.2 subtype, the primary L-type alpha subunit expressed in cardiac myocytes [52].

Together, differentiated SH-SY5Y and differentiated hNPCs share similar properties in gene expression, phenotypes, and responses to oxidative stress, thus, making them suitable model systems for studying the behavior of human dopaminergic neurons. Their usefulness as model systems would be even stronger if it were found that either of these cell lines displayed the characteristic DA neuron pacemaker activity. A limitation of our study is that we haven’t fully characterized the physiological phenotypes of the differentiated cell lines and in particular, we were unable to assess whether the cells’ physiological characteristics are more similar to those of DA neurons found in the ventral tegmental area (VTA) or the substantia nigra (SN) of the brain. This is an important distinction since L-type calcium channels are more involved in the autonomous pacemaker activity of SN DA neurons resulting in greater intracellular Ca^2+^ loads with the associated metabolic costs, and SN DA neurons are more susceptible to degeneration in PD [25, 53]. Although both Cav1.2 and Cav1.3 are expressed in SNc DA neurons [54], the predominant subtype, Cav1.3 L-type channels, which activate at more hyperpolarized potentials, may also be susceptible to oxidation by H_2_O_2_ as they contain cysteine residues (based on our visual inspection of the amino acid sequence of the Cav1.3 α subunit). Determining whether the behavior of the *in-vitro* cell models of human DA neurons is more similar to VTA or SN DA neurons will warrant more investigation to identify which cell models are most appropriate for studying specific brain diseases.

In brain slice preparations, an increase in H_2_O_2_ can result in a hyperpolarization of the membrane potential of SNc DA neurons that is dependent on K_ATP_ channel activity, demonstrating that H_2_O_2_ can also regulate the excitability of DA neurons via the K_ATP_ channels [55]. Our modeling data is not inconsistent with these findings showing hyperpolarization. This hyperpolarization and termination of spontaneous firing activities occur several minutes (4-5 minutes) following the exogenous application of H_2_O_2_ (1.5mM) [55]. Thus, it is plausible that H_2_O_2_-induced hyperpolarization may be a result of a secondary effect rather than a direct effect of H_2_O_2_ on K_ATP_ channels. As ADP accumulates inside the cell due to Ca^2+^-ATPase activity, the buildup in ADP then activates K_ATP_ channels, which causes hyperpolarization. In support of this notion is a study that shows that calcium entry in SN via L-type channels enhances the single-channel opening probability of K_ATP_ [47]. Using computational modeling, they found that calcium entry leads to the accumulation of ADP as a result of Ca-ATPase activity. The activation of K_ATP_ channels in the presence of an NMDA current caused the model to switch from firing single spikes at regular intervals to bursts of spikes. K_ATP_ channels have been shown *in vivo* to regulate burst firing in a subpopulation of SN DA neurons [49]. Our model does not include either K_ATP_ channels or ATP hydrolysis, and so, is unable to predict such secondary effects of H_2_O_2._ It is also not known whether differentiated hNPCs express the K_ATP_ channel gene, KCNJ11. A future extension would be to add these mechanisms as well as synaptic inputs to our model to evaluate whether it is able to better reproduce the calcium oscillations observed in the hNPC cultures.

## CONCLUSIONS

In the human dopaminergic cell lines, SH-SY5Y, and differentiated hNPCS, we found that L-type Ca^2+^ channels were involved in the response to oxidative stress, providing a link between oxidative stress and calcium signaling. Our computational modeling showed that direct potentiation of the L-type channel by exogenous H_2_O_2_, such as that reported in cardiac myocytes, could result in increased internal Ca^2+^ concentration associated with increased oxidative stress and damage to DA neurons.

## ACKNOWLEDGMENTS

We dedicate this manuscript in honor of the late Professor Emeritus Ian M. Cooke. He was a mentor, friend, and colleague to many of us. We thank Dr. Hartline for his leadership in supporting faculty members in the Békésy Lab.

## Abbreviations

Ca2+: calcium
PBS: (Phosphate Buffered Saline)
hNPCs: human neural progenitor cells
DCFH-DA: 2′,7′-dichlorofluorescein diacetate
DHE: dihydroethidium

